# Notch ligand Dll4 impairs cell recruitment into aortic clusters and limits hematopoietic stem cells

**DOI:** 10.1101/2019.12.16.877407

**Authors:** Cristina Porcheri, Ohad Golan, Fernando J. Calero-Nieto, Roshana Thambyrajah, Cristina Ruiz-Herguido, Xiaonan Wang, Francesca Catto, Yolanda Guillen, Roshani Sinha, Jessica González, Sarah J. Kinston, Samanta A. Mariani, Antonio Maglitto, Chris Vink, Elaine Dzierzak, Pierre Charbord, Bertie Göttgens, Lluis Espinosa, David Sprinzak, Anna Bigas

**Author notes:** Corresponding author: Anna Bigas, Cancer Research Program. CIBERONC. Institut Mar d’Investigacions Mèdiques, Doctor Aiguader 88, 08003, Spain. Authors can confirm that all relevant data are included in the paper and/or its supplementary information files. The datasets generated and analysed during the current study are available from the corresponding author on reasonable request. The RNAseq and scRNAseq datasets generated and analysed during the current study (Figure 5 and Supplementary Figure 5) are available in the *GEO* repository, with the accession codes *(submitted)*.

## Abstract

Hematopoietic stem cells (HSCs) develop from the hemogenic endothelium in cluster structures that protrude into the embryonic aortic lumen. Although much is known about the molecular characteristics of the developing hematopoietic cells, we lack a complete understanding of their origin and the three-dimensional organization of the niche. Here we use advanced live imaging techniques of organotypic slice cultures, clonal analysis, and mathematical modelling to show the two-step process of intra-aortic hematopoietic cluster (IACH) formation. First, a hemogenic progenitor buds up from the endothelium and undergoes division forming the monoclonal core of the IAHC. Next, surrounding hemogenic cells are recruited into the IAHC, increasing their size and heterogeneity. We identified the Notch ligand Dll4 as a negative regulator of the recruitment phase of IAHC. Blocking of Dll4 promotes the entrance of new hemogenic Gfi1+ cells into the IAHC and increases the number of cells that acquire HSC activity. Mathematical modelling based on our data provides estimation of the cluster lifetime and the average recruitment time of hemogenic cells to the cluster under physiologic and Dll4-inhibited conditions.

## INTRODUCTION

Hematopoietic stem cells (HSCs) are generated during embryonic life in a process known as endothelial to hematopoietic transition (EHT). EHT takes place in several vessels of the vertebrate embryo and involves the acquisition of hematopoietic features by a specific population of endothelial cells ^1–4^. However, most of the cells that undergo EHT become differentiated hematopoietic cells or progenitors, but do not become HSCs. A number of cells localized in the dorsal aorta in the AGM region (where AGM refers to the Aorta surrounded by Gonads and Mesonephros tissues) will generate HSCs ^5, 6^ and will be the founders of the adult hematopoietic system. Within the AGM region, and in the ventral side of the aorta, a subset of endothelial cells that is characterized by the expression of hemogenic factors such as Gfi1, Runx1 and/or Gata2 are collectively known as hemogenic endothelium. In most vertebrate embryos, functional HSCs develop within hematopoietic clusters that derive from the hemogenic endothelium and then emerge into the lumen of the aorta ^4^. Intra-aortic hematopoietic clusters (IAHC) contain and are the niche of nascent HSCs in most vertebrates, however the mechanisms of IAHC formation and HSC development inside these structures remain elusive. It was previously postulated that bulging of endothelial cells expressing CD41, Kit, Gata2 and/or Gfi1 amongst others, is one of the first events in the formation of hematopoietic clusters ^7–10^. Recent reports studying clonality of cells in the IAHC have determined that they are polyclonal in origin suggesting that at least two independently generated cells aggregate to form a cluster ^5, 11^. In addition, cells in the IAHC are cycling and highly proliferative ^11, 12^.

Beyond these descriptive analyses, the events, mechanisms, and dynamics that determine the architecture of the clusters have never been addressed. Previous published work on signals required for HSC generation converge on the Notch pathway (reviewed in ^13^). Notch receptors can be activated by different ligands either of the Jagged or Delta family of proteins. In the embryonic aortic endothelium, both types of ligands are expressed in a variety of cells and their interplay seems to be crucial to initiate the arterial and hematopoietic programs) ^14–17^.

Here we show by confocal live imaging, clonal genetic studies, single cell RNA-seq and mathematical modeling that IAHCs are originated as clonal entities by cell division, but soon after the first division event, new neighboring cells are recruited into the cluster. We found that the number of cells inside the clusters is tightly controlled by the Notch ligand Dll4, which restrains the recruitment of new cells in the newly-formed clusters. Blockage of Dll4 results in larger polyclonal clusters and affects the number of functional HSCs generated.

## RESULTS

### Formation of IAHC initiates with cell division

To better understand the formation of IAHC, we imaged the first stages of cluster emergence using a reporter for CD41 (Integrin αIIβ), which is one of the earliest genes expressed in the hematopoietic clusters together with kit (Fig1b)^9^. We crossed membrane tethered CD41:YFP reporter mice^18^ with the H2B-GFP transgenic line^19^ and performed time-lapse imaging on organotypic slice culture of E10.5 double transgenic embryos. We consistently identified single CD41:YFP+ cells arising from the endothelial layer and protruding onto the aortic lumen (Fig1a, Fig1c, Movie 1). In agreement with previous observations ^4^, we only detected individual cells, and not groups of CD41:YFP+ cells bulging towards the lumen of the aorta. Time-lapse analysis revealed that individual bulging CD41:YFP+ cells subsequently undergo a cellular division (Fig1a, Fig1d, Movie2) and both CD41:YFP+ daughter cells remained in intimate contact, likely forming the first unit of the IAHC. Within the IAHC, hematopoietic cells are actively cycling (Fig1e, Fig1f, Suppl Fig1), and incorporate BrdU upon continuous administration, until all cells in the clusters are positive (saturation time: 7-8h, Fig1f). Considering that the percentage of BrdU-incorporating cells does not change between 1 and 4 hours of initial BrdU exposure, we assume that the S-phase length (Ts) lasts 4-5 hours. This allows us to estimate the cell cycle length (Tc) duration in 11-13 hours, as a result of the sum between the saturation time (7-8h) and the estimated Ts (4-5h) ^20^. To confirm that all cells in the IAHC are actively cycling, we analyzed the rate of Ki67-expression to find a high number of Ki67 positive cells in the clusters (87,4% ± 5,2) compared to the (59,3% ± 2,92) of the neighboring endothelial cells (Suppl Fig1a, 1b, 1c and 1d). Time-lapse imaging of mitotic CD41:YFP+ cells and staining for the P-Histone3 mitotic marker confirms the existence of cell division inside IAHCs (Suppl Fig 2, Movie3).

**Figure 1:**
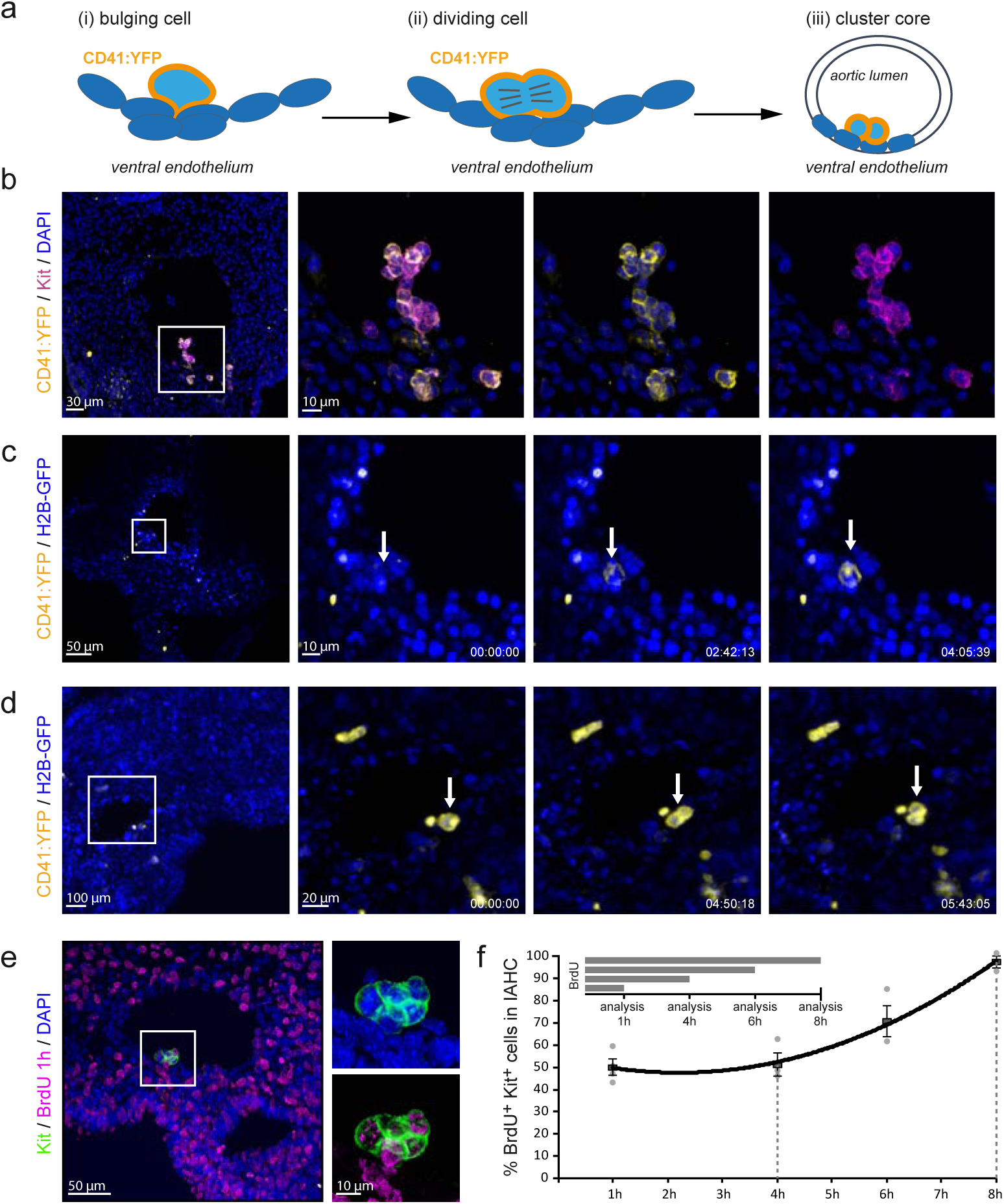
Bulging cells divide to form the monoclonal core of the IAHC. (**a**) Schematic representation of initial events of IAHC formation. (i) A single cell protrudes into the lumen of the aorta and starts expressing CD41. (ii) The CD41:YFP+ bulging cell undergoes mitosis (iii), forming the core of the IAHC. (**b**) Expression of Kit (magenta) largely overlaps with the expression of CD41:YFP (yellow) in IAHC. Multistack reconstruction of confocal images. Scale bars: 30μm overview, 10μm magnification. (**c**) Snapshots of Movie 1. Time-lapse of embryonic organotypic slice culture from CD41:YFP (yellow): H2B-GFP (blue) reporter mouse shows the emergence of a single cell protruding from the endothelial layer into the aortic lumen. Scale bars: 50μm (overview), 10μm (magnification). Time expressed in hh:mm:ss. (**d**) Snapshots of Movie 2. Time-lapse of embryonic organotypic slice culture from CD41:YFP (yellow): H2B-GFP (blue) reporter mouse shows a single CD41:YFP expressing cell that divides once on the aortic endothelium. Scale bars: 100μm (overview), 20μm (magnification). Time expressed in hh:mm:ss. (**e**) BrdU incorporation in the Kit+ IAHC upon cumulative administration. Multistack reconstruction of confocal images. Representative picture of a short BrdU pulse (1h). Scale bars: 50μm (overview), 10μm (magnification). (**f**) Quantification of the percentage of Kit+ cells within IAHCs incorporating BrdU using confocal images. All cells (Labelling Index=100%) can be labelled after 8h, indicating the lack of quiescent cells inside the IAHCs. Graph represents mean±SE.

### IAHCs start monoclonal but become polyclonal after cell division

To test whether hematopoietic clusters were formed by a series of sequential divisions from a single hemogenic precursor, we took advantage of the Confetti reporter mouse crossed with the VE-cadherin-Cre-ERT (VeCadCre^ER^) line. In this model, single pulse administration of Tamoxifen before the time of cluster emergence (E8), allows Cre translocation and recombination of the transgene in cells expressing VE-Cadherin, including the hemogenic endothelial cells that will give rise to clusters (Fig 2a). In our experimental conditions, the overall recombination efficiency of the confetti cassette in the endothelial cells was of 22.92 ± 5.4% (Fig 2b and Fig2c) and allowed the tracking of the sparse single recombined cells. Clusters of monoclonal origin would display one single color (from the originating cell), while in case of a polyclonal origin the cluster will contain cells of various colors or a mix of colored and uncolored cells (Fig 2d, Fig 2e). After labeling cluster cells with the Kit marker, we consistently found that two-cell and three-cell clusters were all single-colored (monoclonal), while clusters with 4 or more cells mostly contained heterogeneous labeling patterns suggestive of a polyclonal origin (Fig 2f and Fig 2g). These results indicated that IAHC formation starts with the generation of a monoclonal core and continues growth with cells from other sources.

**Figure 2:**
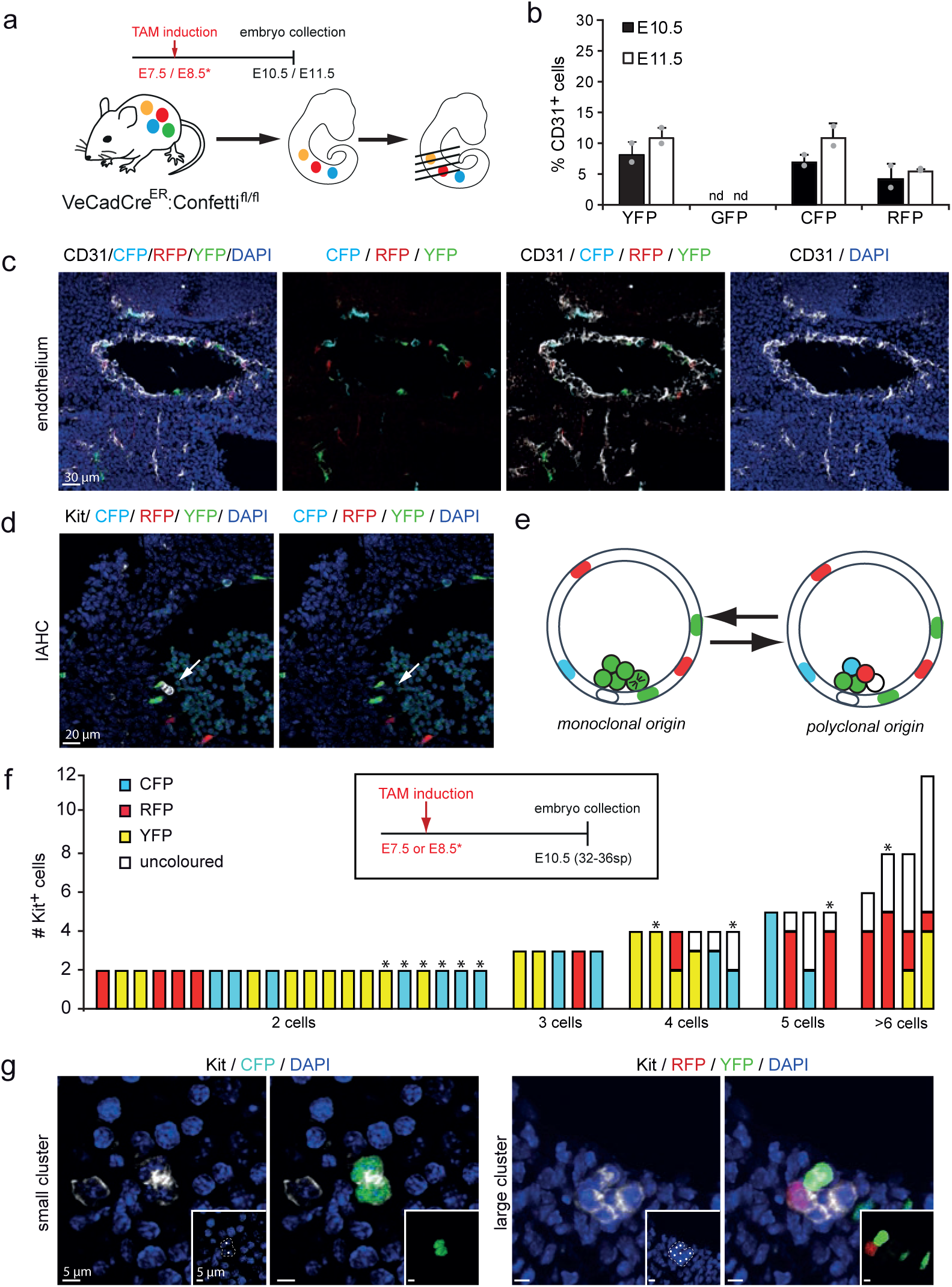
IAHCs have a monoclonal, proliferative origin but become polyclonal at later stage. (**a**) Schematic representation of time of induction of recombination in VeCadCre^ER^:Confetti pregnant females and analysis of embryos. Tamoxifen was injected before cluster initiation (E7.5 or E8.5) and embryos collected at the peak of cluster formation (E10.5 or E11.5). (**b**) Quantification of recombination efficiency at E10.5 and E11.5 using confocal images. Bars represents the mean±SD of the percentage of cells expressing single colored cassettes in CD31+ endothelial cells (n=2). (**c,d**) Transversal section of a recombined VeCadCre^ER^:Confetti AGM from an E10.5 embryo and immunostained with CD31 (**c**) (white in image, black in label) or Kit (**d**) (white in image, black in label). Multistack reconstruction of confocal images. Scale bars 30μm (**c**) and 20 ⎧m (**d**). (**e**) Schematic representation of possible outcomes in the IAHC after recombination: Clusters derived from sequential division of one colored cell will be unicolored (left panel). Clusters will be multicolored (right panel) when derived from several, independently recombined endothelial cells. (**f**) Analyses of recombination in 40 clusters. Bars representing single clusters are grouped based on the number of Kit+ cells. Recombination induction follows the schema in the black square. (n=12 embryo). (**g**) Representative confetti recombination on clusters of different sizes. Small cluster are monoclonal (left panel), while larger clusters are polyclonal (right panel). Inlet shows relevant confetti channel. Kit is shown in white in the image, in black in the label. Multistack reconstruction of confocal images. Scale bars 5 μm.

### The Notch ligand Dll4 is preferentially expressed in small IAHC

Because the Notch pathway is crucial for HSC development ^21, 22^, we performed a detailed study of the expression levels and distribution of Notch elements in the AGM. We found a highly heterogeneous expression of Dll4 ligand in most IAHC and endothelial cells (Fig 3). Confocal microscopy analysis demonstrated that small clusters (2-5 cells) were exclusively formed by Dll4 positive cells, while bigger clusters (>5 cells) contained an increasing proportion of Dll4 negative cells (Fig 3B). In clusters with more than 11 Kit+ cells only 47% ± 10 express the Notch ligand Dll4 (Fig 3A and 3B). In these clusters, 42% showed Dll4+ cells located in the apical portion toward the lumen (Figure 3A), however we could not establish a common pattern since the rest of the analyzed IAHC had Dll4+ cells randomly distributed between basal (endothelial) and apical portion (lumen). In contrast, the Notch ligand Jag1, which is required for downregulation of the endothelial program ^23^ and Notch-dependent activation of hematopoietic genes in the nascent HSCs ^22^ was found preferentially expressed in the endothelial layer of the AGM (Suppl Fig 3). These results suggest that Dll4 is not only required to promote the arterial fate as previously demonstrated, but also plays a role in IAHC formation.

**Figure 3:**
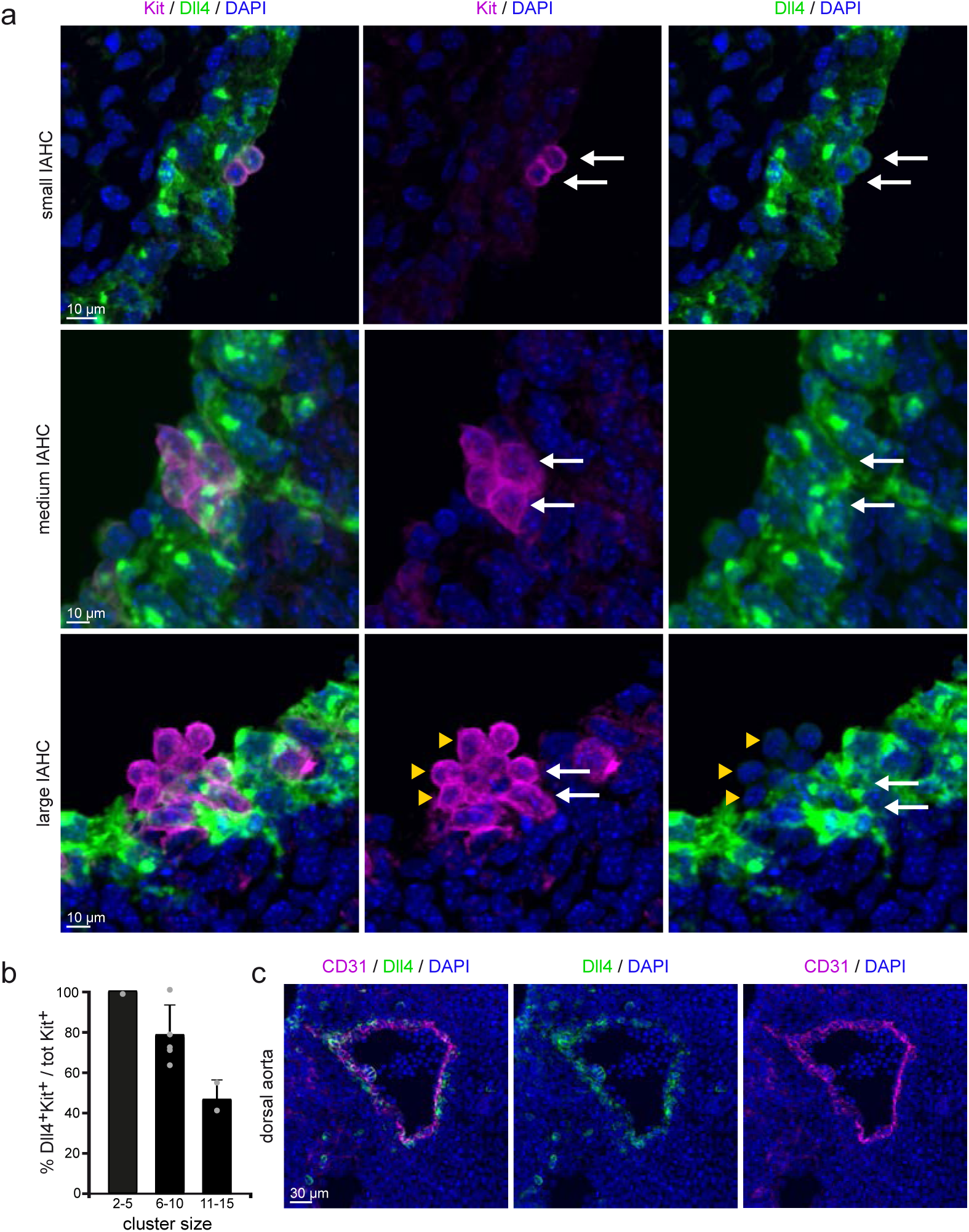
Dll4-ligand expression in the IAHC varies with cluster size. (**a**) Detail images of Dll4 staining in Kit^+^ cells of IAHC of different sizes. Dll4 expression inversely correlates with the size of the cluster: 2-5 cell IAHCs are uniformly Dll4-positive (upper and middle panel), >5 cell IAHCs are partially Dll4 positive (lower panel). Multistack reconstruction of confocal images. Scale bars: 10 μm. (**b**) Quantification of Kit^+^ Dll4^+^ cells using confocal images. Bars represent the percentage of Dll4 expressing cells within a cluster in each category of cluster size. Mean±SD (total 17 clusters analysed, n=2). (**c**) Overview of Dll4-ligand expression in the embryonic aorta at E10.5. Dll4 is expressed in the majority of the endothelial layer. Scale bar: 30μm

### Specific blockage of Dll4 results in increased IAHC size

To investigate the functional contribution of Dll4 in the IAHCs, we used an αDll4 antibody specifically developed to interfere with the Dll4-dependent activation of Notch ^24^. We injected αDll4 or IgG control into the beating heart of the E10-E10.5 embryos (32-37 sp) to minimize the potential side effects of a systemic administration. Following this procedure, we found that the antibody was distributed according to physiologic blood flow and specifically recognized the Dll4-ligand exposed to the aortic lumen (Suppl Fig 4). We then maintained the injected embryos in organotypic culture conditions for 5 hours and processed them for subsequent analyses. Whole mount immunofluorescence of CD31 and Kit staining revealed that the structure of the aorta was preserved after the incubation with αDll4 comparable to IgG control injected embryos (Fig 4a). Kit+ IAHC were similarly distributed along the aorta of control and αDll4 treated embryos while larger IAHCs were found in the latter (Fig 4a, Fig 4b). We then analyzed the size of each cluster per embryo based on the number of Kit+ cells forming them (Fig 4b, Fig 4c, Fig 4d). In control-treated embryos, the majority of IAHC were small (in the range of 2 to 5 cells: 80 ± 3%) and no cluster above 18 cells was detected. In contrast, in the αDll4-treated embryos, the percentage of small IAHC decreased (53. ± 8 %) while bigger clusters were more frequently found, with very large IAHCs (>16 cell) representing 5 ± 3% of total clusters (Fig 4c, 4d). Interestingly, the total number of IAHCs (Fig 4e) or the single kit^+^ cells bulging from the endothelium (Fig 4f) did not change upon αDll4 treatment, suggesting that Dll4 function was confined to the already initiated IAHCs.

**Figure 4:**
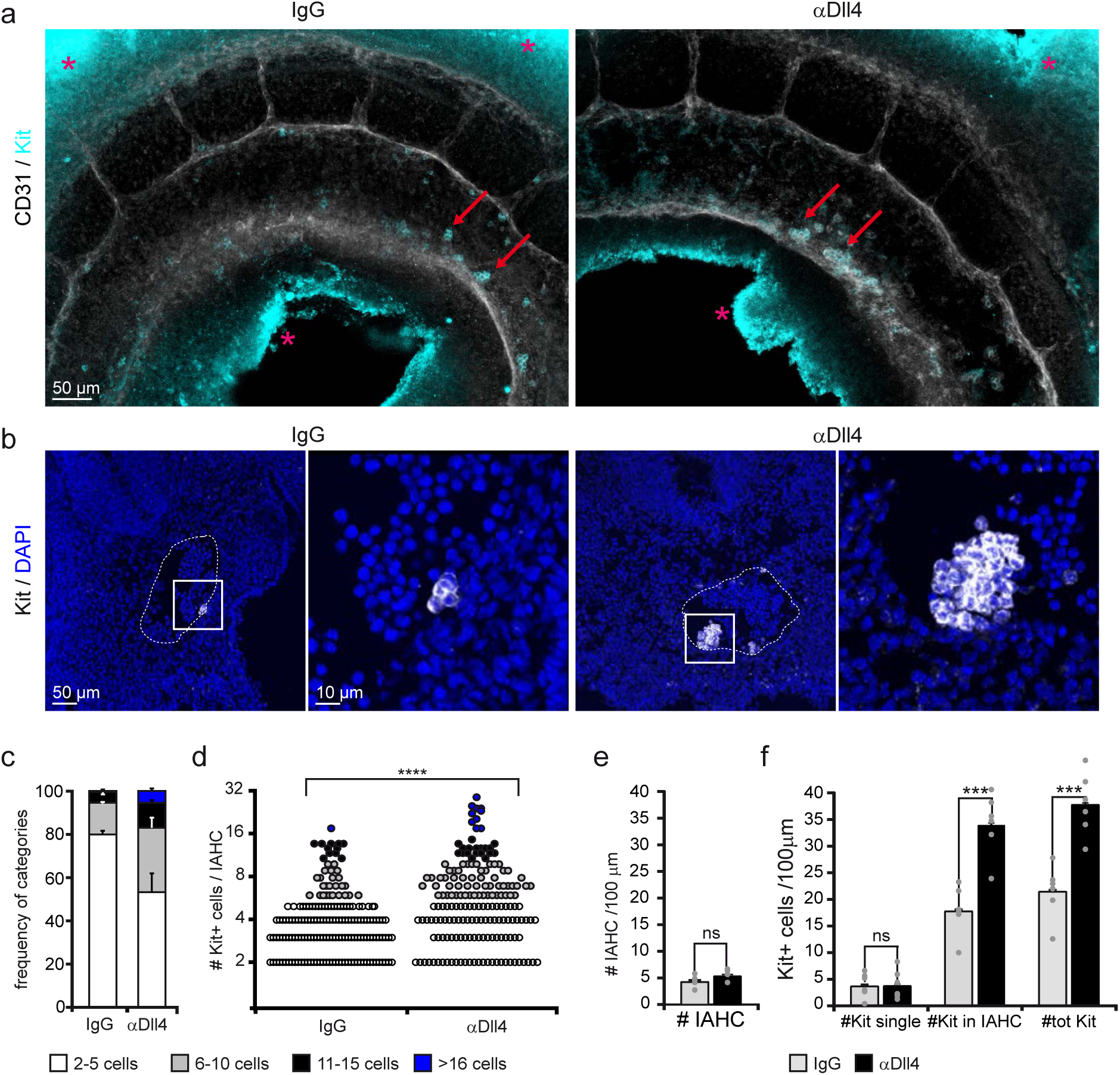
Blockage of Dll4 increases the number of cells in IAHCs. (**a**)Whole mount images of the aorta endothelium stained for CD31 (white in image, black in label) and Kit (IAHC, cyan) upon IgG (left) or αDll4 antibody treatment (right). Multistack reconstruction of confocal images. Red arrows indicate IAHCs on the ventral part of the aortic wall. Asterisks indicate aspecific autofluorescence. Scale bar: 50μm. (**b**) Representative IAHCs of IgG and αDll4 treated embryos. Kit is shown in white in the image, in black in the label. Multistack reconstruction of confocal images. Note the increased IAHC size in the right panel. Scale bars: 50μm (overview), and 10μm (magnification). (**c**) Frequency of IAHC categories based on the number of Kit^+^ cells forming a cluster. Quantification of confocal images. Bars represent the mean±SE (n=6) (**d**) Number of Kit+ cells per IAHC increases following αDll4 antibody treatment. Each dot represents one cluster. Quantification on confocal images. Statistical analysis: Mann–Whitney U test. ****p< 0.0001. (**e**) Total number of IAHCs per 100μm. Quantification of confocal images. Mean±SE. Statistical analysis: t-test. ns p> 0.05; (n=6). (**f**) Quantification of the number of Kit+ cells per 100 μm using confocal images. Bars represent the mean±SE. Statistical analysis: t-test. ns p> 0.05; ***p< 0.001 (n=6).

### Multiple signaling pathways including Notch are altered in Dll4-blocked Kit+CD45-cells

To investigate the mechanism behind the increased cellularity of IAHC after Dll4 blockage, we purified IAHC subpopulations based on kit expression. In particular, we collected 100-300 cells from the Kit+CD45-(containing Pre-HSCs), Kit+CD41+CD45- and Kit+CD45+ (containing HSCs) from 4 different IgG and αDll4 treated embryos and performed RNA-seq (Fig 5a). Differentially expressed genes (DEGs) among the three different populations were assessed by pairwise comparisons, resulting in three sets of up- and down-regulated genes. Intersection of the gene sets allowed discriminating DEGs according to the treatment that were exclusive to each cell population. Principal Component Analysis (PCA) using the total number of DEGs indicated that the samples clearly segregated according to phenotype (Fig 5B). In a second comparison we contrasted control vs αDll4 treated cells and performed a PCA using the DEGs that drove a clear-cut discrimination between treated and untreated samples (Fig 5C). Moreover, both PCA plots revealed that the Kit+CD45-pre-HSC-containing population showed the clearest clusterization based on the treatment. A functional enrichment analysis on the DEGs according to the treatment is shown in Fig 5D. Within the biological processes enriched in dowregulated genes after αDll4 treatment, we found the Notch signaling pathway. In addition to Notch, crucial pathways for HSC development such as NFκB, Wnt and TGFβ pathways were specifically enriched after the treatment, as well as alterations in categories associated with cell remodeling and movement (Fig 5d). Among up-regulated genes, there were genes involved in RNA processing and DNA repair.

**Figure 5:**
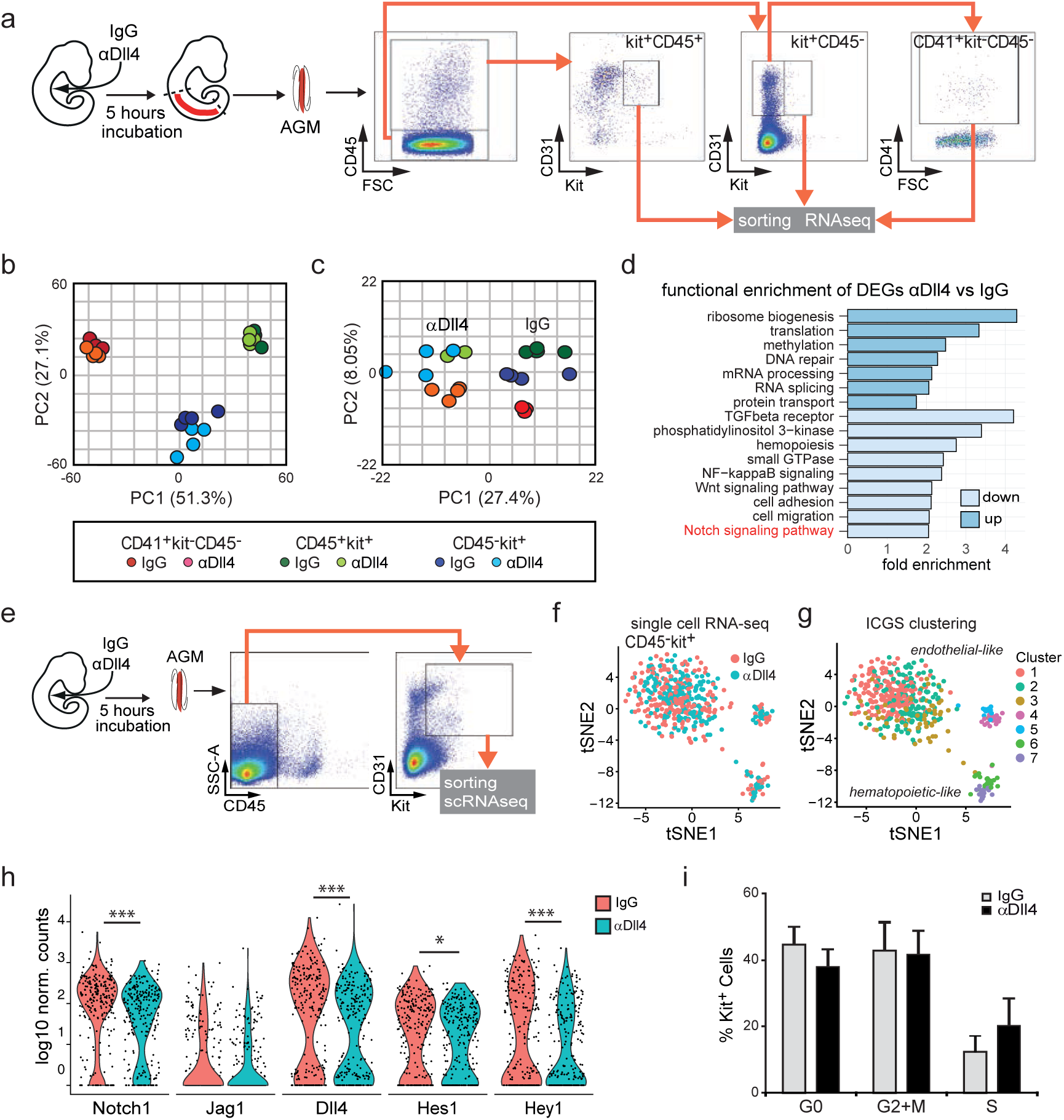
changes in gene expression and decrease in Notch activity after Dll4 blockage. (**a**) Experimental design: E10.5 embryos were injected with IgG or αDll4, incubated for 5h and then dissected. The indicated IAHC subpopulations (kit+CD45+; kit+CD45-;kit-CD45-CD41+ were purified and RNA was sequenced. (**b**) PCA of the different subpopulations from IgG or αDll4 treated embryos based on normalized gene counts.(**c**) PCA considering the DEG according to the treatment specific for each subpopulation. (**d**) Function enrichment analysis of DEG in the Kit+CD45-population according to the treatment(**e**) Experimental design and sorting gate for Kit+CD45-cells for single cell RNAseq (n=483 cells).(**f-g**) tSNE distribution of IgG and αDll4-treated Kit+CD45-cells from single cell RNA-seq data. (**f**) cells corresponding to each treatment are indicated and (**g**) different clusters identified by ICGS (Suppl Fig 5) are represented.1-3 are endothelial-like cells and 6-7 are hematopoietic-like cells.(**h**) Violin plots of Notch family elements expression in IgG and αDll4 treated cells (adj p val: *<0.1 and *** p<0.00001). (**i**) Percentage of cells in each cell cycle phase extracted from single cell RNA-seq analysis. Bars represent the average±SE (n=483).

To further investigate the effect of Dll4 intervention in individual Kit+ precursors, we performed single cell RNA-seq of CD31+kit+CD45-cells (Fig 5e). We analyzed 239 control cells and 244 αDll4 -treated cells from 4 different embryos each (4 untreated (32-34sp) and 4 treated (32-35sp)). Three subpopulations were clearly separated by tSNE, which correlated with 7 cell clusters generated by Iterative Clustering and Guide-gene Selection (ICGS, ^25^) (Fig 5g and Suppl Fig 5a and Suppl Table 1). Based on marker gene expression, the largest cell population corresponded to endothelium while hematopoietic marker gene expression was more prominent in the smaller population. Importantly, integrated analysis with scRNA-Seq from FACS-sorted AGM populations ^26^ confirmed the endothelial and hematopoietic nature of these populations (Suppl. Fig 5b and Suppl Table 1). While there was no readily identifiable shift in cell populations after blocking Dll4, differential gene expression between IgG and αDll4-treated Kit+ cells (Fig 5f) revealed downregulation of several genes associated with the Notch pathway (Fig 5h) and other signaling pathways (Suppl Table 2). Importantly the significance of this observation was confirmed by gene pathway enrichment analysis (Suppl. Fig 5c). Of note, computational analysis of single cell transcriptomes revealed no change in the number of cells in different phases of the cell cycle between both conditions indicating that the number of IAHC cells did not increase by proliferation upon Dll4 blockage (Fig 5h). This conclusion was experimentally supported by demonstrating similar levels of BrdU incorporation after IgG and αDll4 treatment (Suppl Fig 6a).

**Figure 6:**
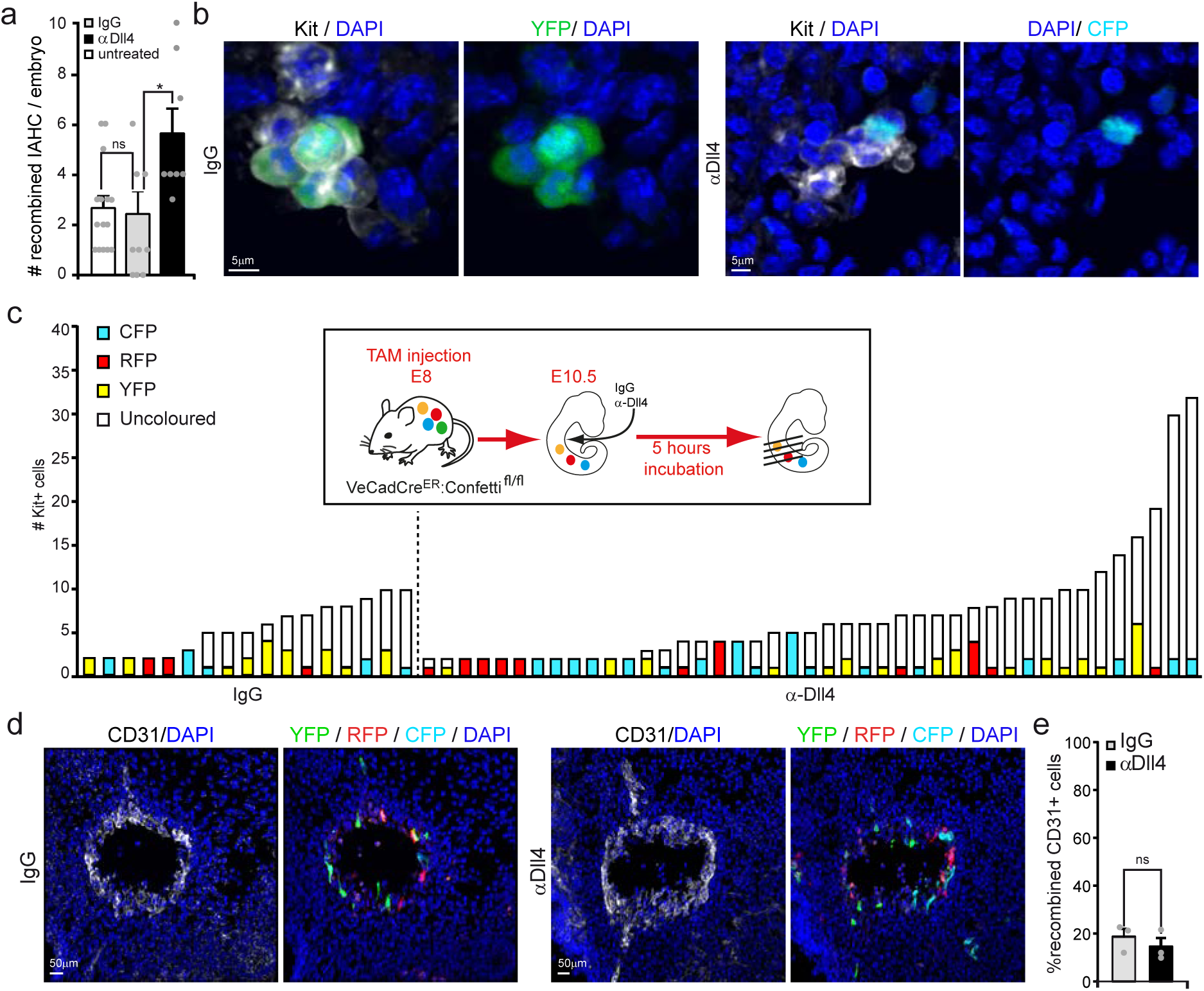
Cell recruitment is increased in Dll4-blocked IAHCs. (**a**) Bar graph represents the frequency of IAHCs containing recombined cells per embryo under the indicated conditions. Quantification of confocal images. Mean±SE( n=15 for untreated, n=7 for IgG and αDll4 treated). (**b**) Representative images of recombined IAHCs following IgG or αDll4 conditions. Kit is shown in white in the image, in black in the label. Multistack reconstruction of confocal images. Scale bar: 5μm. (**c**) Quantification and classification of IAHC containing recombined cells in TAM-induced VeCadCre^ER^:Confetti embryos after IgG or αDll4 treatment. Embryos treated with αDll4 contain more IAHC with colored cells. Quantification of confocal images (n=7 embryos). (**d**) Representative images of recombined endothelial cells in transversal sections of the aorta. The distribution and amount of recombined endothelial cells is similar in IgG and αDll4 conditions. CD31 is shown in white in the image, in black in the label. Multistack reconstruction of confocal images. Scale bar: 50μm. **(e**) Quantification of recombined cells in the aortic endothelium using confocal images. Mean±SE (n=3).

### Recruitment of new cells into the IAHC is controlled by Dll4

Having established that increased proliferation is unlikely to account for increased cluster sizes, our second hypothesis was that IAHC grow by recruitment of new cells into the already initiated IAHC. To test whether Dll4 regulates the capacity of endothelial cells to be recruited into the nascent IACH, we used TAM-induced VeCadCre^ER^:confetti embryos (Fig 6). We intracardiacally injected IgG or αDll4 followed by 5 hours incubation and processed the embryos for Kit staining (n=7 per condition). We confirmed that transgenic modifications per se did not alter the total number nor the size of IAHCs and bigger clusters formed as a consequence of αDll4 blockage (Suppl Fig 7).

In these experiments, we observed more IAHC with colored cells in the αDll4-treated condition compared to IgG (Fig 6a; Fig 6c). This increased frequency is not associated with enhanced recombination in the endothelium (Fig 6d; Fig 6e) nor an increase in the total number of IAHC formed (Suppl Fig 7). Our interpretation is that recruitment events occur more often upon αDll4 treatment, leading to more IAHCs incorporating a colored cell. Consistently, the number of IAHC containing recombined cells per embryo is increased in these conditions (Fig6a; Fig6c). Collectively, these data indicate that αDll4 treatment increases the number of cells in IAHC by facilitating the chance of a colored cell to be incorporated in the growing cluster.

### Mathematical model supports a role for Dll4 controlling cell recruitment in IAHC

To better understand the parameters that control IAHC formation, we developed a probabilistic simulation of the process. In particular, we wanted to check what are the recruitment rates and cluster life-times that can explain the measured distributions of colored vs non-colored cells in IAHCs both in the IgG and αDll4 treated samples. The model assumes that clusters can be initiated with the same probability at any time point and can be detached after a certain time (defined as cluster life time). Since in physiological condition two-cell clusters are either all colored or all non-colored, we assume that the initiation stage always starts with two cells. Clusters in the model can then grow either by cell division events (assumed to happen every 11-13 hours, based on the BrdU analysis in Fig. 1F), or by recruitment events (see schematic in Fig. 7a). We focused on determining two parameters: (1) cluster life time and (2) average recruitment time (or recruitment probability). To determine the values of the parameters that best fit the measured IgG treated distribution, we performed 4000 simulations for each parameter set and checked whether the resulting distribution fit the experimental cluster sizes, and the fraction of colored cells in clusters (see details of the simulation in the methods). We found that values that best fit the measured IgG treated distribution are obtained for cluster life-time of 21 hours (90% confidence interval (CI) 10-22 hours) and average recruitment time is 11 hours (90% CI 3-18 hours) (Fig. 7b). We next simulated the case where samples were treated with αDll4 for 5 hours. Here, we set the cluster life time to 21 hours (as obtained from the analysis of the IgG treated clusters) and checked what is the average recruitment time that can fit the observed data. We find that the best fit is obtained for recruitment time of 5 hours (90% CI 3-8 hours), which is more than twice faster than the recruitment time without blocking Dll4 (Fig. 7c). Using these parameters we run a simulation test for IgG and αDll4 treatment conditions, which resulted in a distribution of cluster size very similar to the one previously observed using empirical data (Fig 7d compared to Fig 6c).

There are several deviations between the cluster distributions observed in the simplified model proposed and the observed cluster distributions, potentially revealing additional processes affecting cluster sizes. First, some of the clusters observed in the αDll4 treated samples have more than 20 cells, significantly above the maximal cluster size obtained in our simulation. It is likely that such clusters arise due to effective merger of nearby clusters.

**Figure 7:**
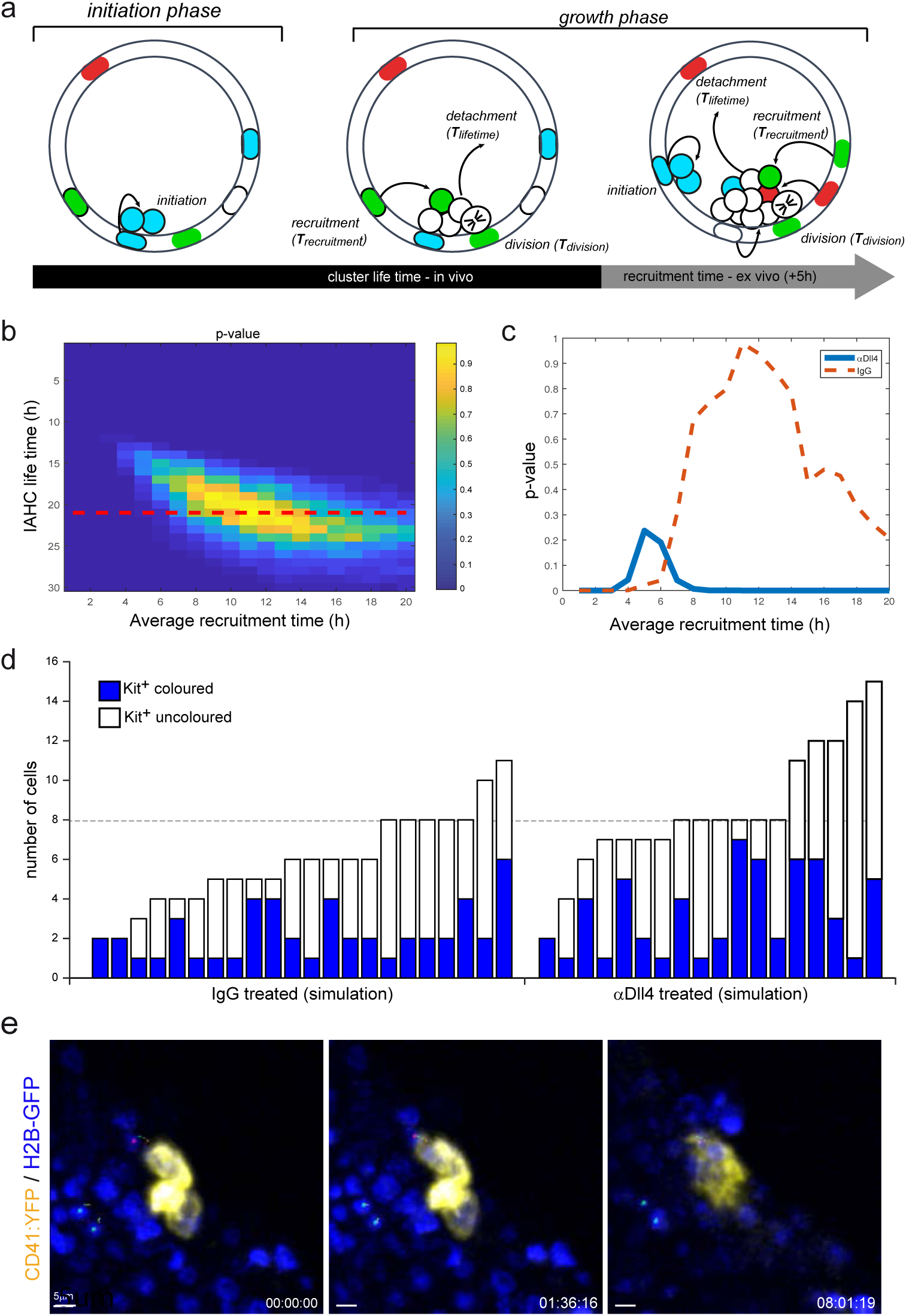
Mathematical model supports a role for Dll4 in controlling cell recruitment into IAHCs. (**a**) Schematics of the probabilistic model of IAHC formation. The processes considered in the model are (1) initiation of cluster, (2) cell division event, (3) recruitment event, and (4) cluster detachment. Each process in the model is described as a typical time scale (*T*_*cluster lifetime*_, *T*_*cell division*_, *T*_*recruitment*_). In case of αDll4 treatment, the final five hours of the simulation are assumed to have a different recruitment time. See details of the simulation in the methods. (**b**) Heat map showing the probability (or p-value) that the simulation produces mean cluster size and fraction of colored cells that match those observed experimentally for the IgG treatment. This probability is calculated for different values of cluster lifetimes and mean recruitment times (values in x and y). 4000 simulations were performed for each set of parameters (each point in the heat map). The most likely parameter values (yellow) correspond to a cluster lifetime of 21 hours and mean recruitment time of 11 hours. Red dashed line indicates the cross-section with the most likely cluster lifetime (21 hours). (**c**) A graph showing the probability that the simulation produces mean cluster size and fraction of colored cells that match those observed experimentally for the αDll4 treated clusters (blue line). Here, the cluster life time is taken to be 21 hours (taken from b) and the probability to match the experimental data is calculated for different recruitment times in the final 5 hours of the simulation (x-axis). The most likely mean recruitment time for the αDll4 treatment is 5 hours (peak of blue line). For comparison, the most likely mean recruitment time for the IgG treatment is 11 hours (Red dashed line, corresponding to the red dashed in (b)). Hotelling’s T-Squared test. (**d**) Examples of simulations of IAHC distributions for the IgG (left) or αDll4 (right) treated samples. Simulations were performed with the most likely parameter values described in b and c. Note that only clusters that have at least one colored cell are shown (i.e clusters with non-colored cells are not represented).(**e**) Snapshots of Movie 5. Time Lapse of embryonic organotypic slice from a CD41:YFP: H2B-GFP reporter mouse showing the recruitment of CD41:YFP negative cells (magenta central point-tracking analyses) migrating towards the IAHC (CD41:YFP+ cells). Green central point and relative tracking identify cells with non-directional movement. Scale bar: 5μm. Time expressed in hh:mm:ss.

Simulations performed assuming that the bigger clusters (>20 cells) are a result of a merger of two smaller clusters with half the size each, resulted in a similar estimation of the recruitment times (Suppl Fig. 8). Second, we do observe two two-cell clusters with one colored cell in the αDll4 treated samples. This may suggest that recruitment times at the two-cell stage may be more significantly affected by the αDll4 treatment. Additional experiments may be required to resolve the recruitment times at different stages.

To further support the conclusion of the model that the increase in cluster size in αDll4 treated embryos is due to more frequent recruitment events, we imaged CD41:YFP^+^;H2B-GFP embryo organotypic slices. In these conditions we indeed identified cells adjacent to a CD41:YFP+ cluster getting incorporated into the cluster and turning on CD41:YFP expression (Fig 7d and Movie 5). We also used organotypic slices from H2B-GFP embryo intracardiacally injected with CD31^549^ antibody, and track the movement of a peripheral cell reaching the center of the close-by cluster (Movie 6). Overall, the confetti analysis, the mathematical modelling and the additional movies provide strong evidence that blocking of Dll4 significantly enhances recruitment of cells into the IAHC.

### Recruitment of hemogenic Gfi^+^ cells into the IAHC and progression to HSCs is restricted by Dll4

To functionally validate the effect of blocking Dll4 in the hematopoietic activity of the IAHC cells, we performed 5h ex-vivo cultures of αDll4 treated and control AGM to test them for HSC and progenitor activity (Fig 8a). We dissected AGMs from IgG or αDll4 injected embryos, and after 48 hours we analyzed CFC activity (Fig 8B). We did not observe major changes in the number and quality of CFC, but a slight increase in the CFC-GM category after Dll4 blockage (Fig 8b). Next, we tested whether αDll4 treatment was directly affecting the number or quality of hemogenic cells. To identify hemogenic cells, we took advantage of the Gfi1:tomato reporter mouse model which labels early hemogenic cells that are not yet expressing Kit or CD45. We found that Gfi1+ cells (hemogenic endothelial precursors) from early embryos (30-32sp) were especially responsive to Dll4 blockage as they significantly transformed into Kit+CD45+ cells when compared with control-treated littermates after 5 hours treatment (Fig 8c, 8d and 8e). These results indicate that Dll4 signaling restricts the incorporation of hemogenic cells into the IAHCs.

**Figure 8:**
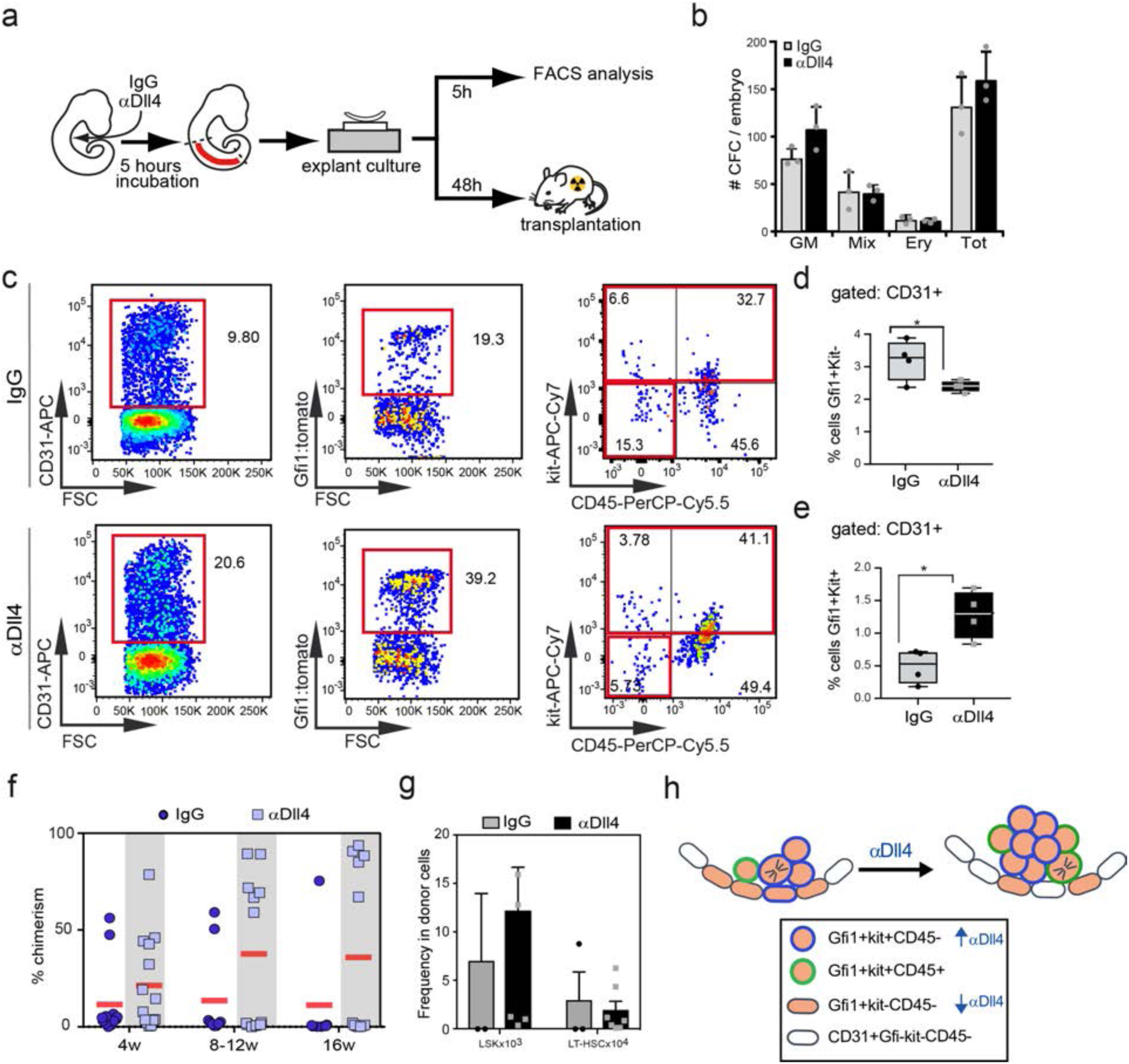
Interference with the Dll4 ligand functionality results in increased hematopoietic potential in embryonic AGM. (**a**) Schematic representation of functional assays for testing the hematopoietic potential of AGM. Embryos were injected intracardiacally and incubated for five hours with αDll4 or IgG control. Explant cultures were either exposed to the relevant treatment for 5h before FACS analyses or incubated for 48h before transplantation into irradiated mice. (**b**) Colony-forming assays using E10.5 AGMs derived from embryos treated with IgG or αDll4. Mean±SD (n=3) No significant differences were detected (**c-e**) FACS analyses of Gfi1^+^ cells derived from explants treated with IgG or αDll4 for 5 hours (**c**). Analyses of the hemogenic Gfi1^+^Kit^-^(**d**) and the Gfi1^+^Kit^+^ (**e**) within the CD31^+^CD45^-^ population. Statistical analysis: t-test. Box plots show upper and lower quartile; center line represents mean (n=4). *p< 0.05; **p< 0.005; non significant comparisons are not indicated (**f**) Percentage of reconstitution (chimerism determined by CD45.1 and CD45.2 expression) in peripheral blood (PB). Embryos treated with IgG or αDll4 were kept in explant culture for 48hours and then transplanted into irradiated recipients. Peripheral blood was analyzed at different time points. Each dot/square represents a single transplanted mouse. (**g**) Analysis of progenitors (lin-sca1+kit+) and HSCs (Lin-sca1+kit+CD48-CD150+) in the donor population in the bone marrow after 4 months of transplantation. (**h**) Model for recruitment of Gfi1^+^ cells into the IAHC. Recruitment of Gfi1^+^endothelial cells increases upon Dll4 blockage.

Finally, we performed transplantation experiments to determine the effect of αDll4 treatment in HSC activity. AGM explants obtained from 30-34 sp CD45.2 embryos were injected with IgG or Dll4 blocker and then processed for hematopoietic transplantation together with 50,000 bone marrow CD45.1/2 support cells into CD45.1 lethally irradiated recipients (Fig 8a). Mice were monthly analysed for the presence of donor CD45.2 cells in the peripheral blood and 4 months after transplantation they were euthanized. We found that 5 of 12 mice transplanted with αDll4 treated AGM cells were highly reconstituted compared with only 2 of 8 mice transplanted with IgG control AGM (Fig 8f). Donor cells contributed to all different lineages in a similar way, but a higher contribution to the B-cell lineage from the αDll4-treated cells was observed (Suppl Fig 9). Importantly, Lin-sca1+Kit+ (LSK) progenitors and CD150+CD48-HSCs were similarly preserved in the αDll4-treated engraftments (Fig 8g). Altogether these data strongly suggest that αDll4-treatment enhances the presence of engraftable LT-HSC without compromising their stemness.

In summary, our results indicate that at initial stages of IAHC formation, Dll4 is restricting the recruitment of Gfi1+ hemogenic cells into the cluster. Once the cluster reaches three-four cells size, Dll4 is decreased and new Gfi1+ hemogenic cells can move into the IAHC (model in Fig 8h).

## DISCUSSION

In this study, through a detailed characterization of the earliest stages of hematopoietic cluster formation, we identify a new role for the Notch-Dll4 axis in regulating the origin of HSC. The emergence of the first definitive HSCs is a complex process requiring formation of specialized structures in the AGM that function as a supportive environment. During mammalian development, the appearance of HSCs correlates with the formation of IAHCs, which expose progenitors to the required set of molecular signaling for their specification and maturation. Despite the fact that IAHCs play an essential role in early hematopoiesis of higher vertebrates (reptiles, avian and mammals), little is known on how they generate and develop. Here, we describe for the first time, the two-step process of the formation of hematopoietic clusters, with an initial monoclonal phase as a result of one or two cell divisions, followed by a recruitment phase that contributes -together with proliferation-to cluster growth. We now provide data on cell cycle kinetics of the cluster that agrees with previous reports^12^ but the focus of the current investigations is on the impact of proliferation on IAHC clonality and architecture. Although polyclonality of IAHC have been suggested ^5, 11^, we now provide evidence for an initial monoclonal origin and the mechanism on how the polyclonality of larger IAHC is achieved and regulated by Notch-Dll4 signals.

The Notch ligand Dll4 has a crucial role in arterial development, which limited genetic studies on aortic hematopoiesis due to the lack of specificity in targeting cluster cells. We now found that Dll4 is expressed in the whole IAHC in small clusters (2-4 cells), but its expression is restricted to few cells in larger IAHC. This pattern suggests a defined function at different phases of IAHC growth that was addressed by testing the effect of αDll4 blocking antibodies in the embryonic aorta. When Dll4 is blocked, IAHC size increases dramatically from newly recruited neighboring cells, as shown by three independent approaches: Imaging of the confetti system shows increased heterogeneity in the mature clusters, time-lapse analyses of CD41:YFP^+^ cells being incorporated into the IAHC and changes in the percentage of Gfi1+ hemogenic cells. These results indicate that Dll4 might block the recruitment events until the IAHC core is completed. Once the initial cluster unit has been formed, few cells in the IAHC downregulate their levels of Dll4 and the break is lifted, allowing recruitment to happen and consequently the cluster to grow. Our results show that the total number of IAHC is not increased after Dll4 blockage, indicating that the emergence of new IAHC is not dependent on Dll4-signalling. Instead, cells in already initiated IAHC (2-4 cells) express high levels of Dll4 and prevent the entrance of new cells until Dll4 is downregulated.

The mechanism of cell recruitment in IAHC is reminiscent of the tip-stalk cell communication in angiogenic sprouting. Tip cells contain high Dll4 that signals to activate Notch in stalk cells and maintain the stalk phenotype. Jagged1 is present in the stalk cells and antagonizes Dll4 activation of Notch1 ^27^ while Fringe will potentiate the action of Dll4. In the IAHC, Dll4 is expressed in small clusters (similar to tip cells) and should signal to the endothelial cells preventing their inclusion into the cluster (similar to stalk cells). When Jag1 counteracts Dll4 signal, either by fringe expression or by cis-inhibition (reviewed in ^28^) Notch signal is inhibited and cells can move into the cluster. The role of fringe in cluster formation has not been explored yet, although expression of Manic fringe is observed in RNA-seq and immunostaining experiments (data not shown). The presence of fringe may inhibit Notch activation by Jag1 and allow Dll4 signaling until the cells reach the cluster.

The specific function(s) of Notch in HSC development is not well understood. While reported data agrees that the level of Notch activity needs to be lower or inhibited from the endothelial cells to generate HSCs ^29–33^, it is also well established that transcriptional Notch activity is required for this process ^22, 32–34^. However, the target cell(s) for Notch activation is unknown. We now show that hemogenic cells (Gfi1+Kit-CD41-CD45-) are one of the targets for Dll4 signal while maintained in the endothelium. When Dll4 is reduced, cells can proceed to the IAHC where HSC can be generated. Our interpretation is that several Notch signals are required for HSC development, likely coming from different ligands (Jag1 and Dll4) ^23^, but also from different Notch receptors (Notch1 and Notch2)^14^.

Experiments using the Gfi1^+^ reporters support that cells recruited to the IAHC are hemogenic. The expression of Gfi1 identifies the first events of hemogenic conversion before the expression of Kit and CD45 which will be acquired in the HSC population ^10^. Our results show that αDll4 treatment decreases the number of Gfi1^+^Kit-hemogenic cells and induces the formation of Gfi1+Kit+. This conversion is very robust when using 30-32 somite pair embryos strongly suggesting that this mechanism is crucial in the initial phases of definitive HSC development. Moreover, functional experiments testing HSC activity indicate that blocking αDll4 antibodies increase the engraftment potential of the early hemogenic cell, additionally suggesting that IAHC of bigger size contain the most mature HSC. Multilineage engraftment is observed from the αDll4-treated HSCs, although we find a higher potential for B-cell repopulation and surprisingly T-cell potential is unaffected, which will be a future matter of investigation. Single cell RNA sequencing data also confirms that the nature of Kit+ cells is unchanged at a transcriptomic level and altogether indicates that blocking Dll4 does not disturb cell fate of HSCs, but it favors its generation or maturation.

Based on our imaging and clonality data, we developed a probabilistic simulation of IAHC formation. The observed distribution of colored and non-colored cells in IAHCs are captured in the model by assuming finite cluster life time of ∼21 hours and an average recruitment time of ∼11 hours. Furthermore, the observed distributions in the αDll4 treated embryo can be best fitted to a recruitment time of only ∼5 hours. We do note that we have observed a few clusters in the αDll4 treated embryos with >30 cells. Such large clusters are unlikely to arise from single recruitment events and indeed do not occur in the simulation. It may be possible that such large clusters emerge from fusion of smaller clusters. Hence, our quantitative modeling approach can test whether the observed IAHC formation can be explained by a combination of cell divisions, recruitment events and cluster detachments, and provide insights into the specific timescales associated with these processes. Moreover, our model can be used to determine the effect of physiological signals (e.g. Notch signaling) on these rates and hence helps understanding the molecular factors that affect HSC generation.

## MATERIAL AND METHODS

### Animals

C57BL/6 J wild-type, (Charles River Laboratories), CD41:YFP ^tg/tg^ ^18^,H2B-GFP ^tg/tg^ ^19^ VeCadCre^ERT35^ and R26R-Confetti ^36^; Gfi1:tomato ^10^ transgenic lines were used. Animals were kept under pathogen-free conditions, and all procedures were approved by the Animal Care Committee of the Parc de Recerca Biomedica de Barcelona, Licence number 9309 approved by the Generalitat de Catalunya. Embryos were obtained from timed pregnant females and staged by somite counting: E10.5 (32–38 sp). The detection of the vaginal plug was designated as day 0.5. Induction of recombination for the VeCadherinCre^ERT^:R26R-Confetti line was performed injecting 50mg/Kg Tamoxifen (Sigma) into the pregnant female at E7.5 or E8.5. Cumulative intraperitoneal injections of 50mg/Kg BrdU was injected at different time point for kinetic analysis as previously described ^20^. Pregnant females were euthanized after 1h BrdU injection and embryo processed for immunostaining. For cumulative analyses pregnant females received BrdU every 2 hours and then euthanized after 4h, 6h and 8hours for embryo collection and processing.

### Dll4-blocking and embryo culture

C57BL/6 J wild-type or VeCadherinCre^ERT^:R26R-Confetti embryos were collected at E10.5 and placed in sterile PBS 10% inactivated fetal bovine serum (FBS) for placenta, yolk sac and tail removal. Blocking antibody Dll4 (Genentech) or mock antibody IgG (Goat Anti-Human Ig, Southern Biotech 2010-01) at 1 μg ml^−1^ were slowly injected intracardiacally until complete clearance of the aortic blood. Treated embryos were then incubated for additional 5hours at 37 °C in a humidified atmosphere with 5% CO_2_ in complete myeloid long-term medium (STEMCELL Technologies - supplemented with 10 ng ml^−1^ interleukin (IL)-3 and 10 µM hydrocortisone (Sigma-Aldrich) containing 5 μg ml^−1^ of the relevant antibody. For BrdU-incorporation analysis intraperitoneal injections of 50mg/Kg BrdU was injected once to the pregnant female and embryos processed as above.

### Organotypic slice culture and Time-lapse imaging and embryonic culture

CD41:YFP^tg/tg^ crossed with H2B-GFP ^tg/tg^ embryos were collected at E10.5. Embryos were placed in sterile PBS 10% FBS for placenta and yolk sack removal. For direct staining of aortic endothelium 5ul of CD31^594^ directly conjugated antibody (BD Bioscience) was injected into the beating heart and whole embryos were incubated in ice for 20min. Embryos were then sectioned using a MacIllwain tissue chopper to obtain horizontal slices of 100um thickness. Organotypic slices were then transferred onto an O-ring chamber and trapped in Agarose 1% (Sigma-Aldrich). Slices were kept in myeloid long-term medium (STEMCELL Technologies - supplemented with 10 ng ml^−1^ interleukin (IL)-3 and 10 µM hydrocortisone (Sigma-Aldrich) in a humidified chamber with 5% CO_2_ at 37 °C for the whole duration of the live imaging. Time Lapse was performed using an inverted Leica-SP5 confocal with excitation lasers 488, 514 and 549 wavelenght as previously described ^4^.

### Immunostaining

For tissue-section immunostainings, embryos were fixed 1h in 4% paraformaldehyde (Sigma-Aldrich) at 4 °C, included in Optimal Cutting Temperature (OCT) (Tissue-Tek, Sakura) and sectioned at 25 μm. Permeabilization was performed with 0,5% triton for 30min at RT and blocking step was done with a solution of 10% FBS and 0,1%triton for 1hour at RT. Primary antibodies were used at the following concentrations: anti-cKit 1:100 (rat, BD); anti-pH3 1:300 (rabbit, Upstate Cell Signaling); anti-GFP 1:300 (rabbit, Molecular probes); anti-Jag1 1:400 (sc-6011-Santa Cruz), anti-Dll4 1:200 (goat, R&D AF1389), anti-Dll4 1:1000 (human, Genentech) anti-ICN1 1:100 (α-N1Icv monoclonal antibody Cell signaling #4147S); anti-BrdU 1:50 (ms, BD Bioscience); Ki67 1:100 (ms, Novacastra); biotinylated anti-CD31 1:100 (BD 553371).

For mouse antibodies, slices were additionally incubated with FAB 1:10 (Roche) in PBS 2h at RT.

For CD31 staining, an additional blocking of endogenous biotin was performed (Endogenous Avidin+Biotin Blocking System, ab3387 abcam) and detected with streptavidin Alexa Red 555 (BD 32355).

For BrdU and Ki67 staining a primary antigen retrieval was performed in 10 mM sodium citrate pH6, 30 min at 80°C. For BrdU staining an additional incubation in HCl 2N for 20min was performed. Fluorochrome-conjugated (1:1000 Alexa Fluor 488,546,647) or horseradish peroxidase (HRP)-conjugated (1:200, Dako) secondary antibodies were used for detection. Three embryos per groups were analysed at each time point and positivity on total number of 4,6-diamidino-2-phenylindole (DAPI; Invitrogen)-positive nuclei counted inside the clusters.

Whole-mount immunostaining was performed as described ^3^. Briefly, embryos were fixed for 20 min in 2% paraformaldehyde, dehydrated in methanol and trimmed. Primary antibodies were used at the following concentration: anti-cKit 1:100 (rat, BD Bioscience), and biotinylated anti-CD31 1:100 (BD 553371). Detection was achieved with horseradish peroxidase (HRP)-conjugated secondary antibody (1:200, Dako) developed with tyramide amplification system TSA (PerkinElmer) and streptavidin Alexa Red 555 (BD 32355).

To analyze the IAHC at a rostro-caudal level, a range of 250-1500 µm of aortic endothelium was sectioned and immunostained with the kit antibody. Section reconstruction from confocal images was performed and the number of Kit^+^ cells and IAHC was quantified in individual aortas. IAHCs were classified: 2-5 cells small cluster; 6-10 cells medium cluster; 11-15 cells large cluster; above 16 cells very large cluster.

### Equipment and settings

All immunofluorescence images were produced using a Confocal Leica SP5 or SP8 with the following settings: 1024X1024 pixel dimension; 400Hz laser; frame average 3 or 4. photomultipliers PMT or Hyd. Fluorochromes used: Alexa 488,546,594,647 or CFP,RFP,YFP,GFP excitation for Confetti analyses. A range of 25μm (standard) to 70μm (whole mount) thick Z-stack was analyzed collecting images every 2.5 μm (standard) or 10 μm (whole mount). Images were processed using Imaris 7.0 and 8.0 software, Bitplane. Gaussian smoothing was applied to all channels when needed and equally to all images of the same batch. For Time-lapse videos a Leica SP5 inverted confocal microscope equipped with CO2 and temperature controlled chamber (CO2=5%, T=37°C) was used. Laser scanned the samples every 15-19min at 700Hz for a maximal time of 9 hours. Movies were processed post-imaging with ImageJ-Fiji to correct drifting (ImageJ-Drift Correction) and recorded in Imaris 8.0 (Bitplane) for further analysis and automatic tracking. Tracking analyses over time was performed using the spot function and central point automatic tracking in Imaris.

### Haematopoietic progenitor assay

C57BL/6 J wild-type embryos at 32-35sp were collected and intracardiacally injected with control IgG or blocking antibody αDll4 as described above. After 5h incubation AGMs were incubated for 20 min at 37 °C in 0.12% collagenase (Sigma-Aldrich). Single cell suspension was obtained and seeded in quadruplicates in Methocult M-3434 semi-solid medium (Stem Cell Technologies) at 37 °C with 5% CO_2_. Colony-forming units were counted after 7 days (n=3 embryos/group).

### Fluorescence-activated cell sorter (FACS) analysis and purification

AGMs were harvested and dissociated in single-cell suspension. Cells were then washed with PBS+10% FBS before antibody staining. Antibody staining was performed in PBS supplemented with 10% FCS in the dark, at room temperature for 15 min, or carried out on ice for 30 min. CD117 (c-kit; APC-Cy7 or APC-eFluor780), CD31 (APC or FITC), CD45.1 (APC-Cy7) and CD45.2 (FITC), Dll4 (PerCP5 or APC) Jag1 (PE or FITC), Ter119 (Pe-Cy7) Ki67 (APC) antibodies were purchased from BD (Franklin Lakes, NJ, USA). DAPI (4,6-diamidino-2-phenylindole; D1306, Invitrogen, Waltham, MA, USA) was used for viability. FACS was performed on LSRII (BD), sorting on FACSAria (BD) and data were analyzed with FlowJo v.10 (TreeStar, Inc., Ashland, OR, USA).

### AGM explants and Transplantation experiments

Charles River C57BL/6 (CD45.2) WT embryos of 32-34sp were treated as described above injecting intracardiacally αDll4 or anti-IgG. AGM were then dissected and cultured as explant as previously described ^37^. In brief, AGMs were deposited on nylon filters (Millipore) placed on metallic supports and cultured in myeloid long-term culture medium (Stem Cell Technologies) supplemented with 10 µM hydrocortisone (Sigma-Aldrich). After 24hours the explanted AGMs were digested in 0.1% collagenase (Sigma-Aldrich) in PBS supplemented with 10% FBS for 20 min at 37°C and used for cell transplantation. Donor cells (CD45.2) were transplanted together with 200 000 BM support cells (CD45.1) into lethally irradiated (8 Gy) recipients (CD45.1). Peripheral blood (PB) donor chimerism was analyzed by FACS at 4, 10-12 and 16 weeks. Lineage analysis on BM-derived cells was performed at 16 weeks post-transplant by flow cytometry with specific antibodies: CD3, B220, Ter119, Mac1, and Gr1 (mouse lineage panel; BD).

### Bulk RNA sequencing and analysis

For RNA-seq study, the SMARTer UltraLow RNA kit v.4 for Illumina Sequencing was used. Briefly, αDll4 and control-treated AGM were digested with 0.1% collagenase and single-cell suspension was stained as described. Different subpopulations were directly sorted in a total volume of 10 μl of reaction buffer and processed for obtaining cDNA following manufacture’s protocol. cDNA amplification was performed by Long Distance PCR (LD-PCR) and the PCR-amplified cDNA purified by immobilization on AMPure XP beads (Agencourt AMPure XP kit). Samples were analyzed with Agilent High Sensitivity DNA Kit (Agilent, Cat. No. 5067-4626).

Covaris shearing was used for Illumina Low input sample preparation and Double last bead purification was performed to remove fragments below 200 bp. Next, samples were used to generate an Illumina sequencing library by NEBNext Ultra protocol following kit instructions. After PCR amplification of the library, the quality was checked on a Bioanalyzer and total cDNA was sequenced using an Illumina HiSeq 2000 sequencer.

MultiQC tool (Ewels et al. 2016) implementing QualiMAP (Quality control), STAR (RNA-seq aligner) and DESeq2 (gene differential expression) packages, was run to obtain normalized gene counts and asses differential gene expression levels among three cell populations Kp45p, Kp45m and Km41p. To test for statistically significant differences between populations, a 1-way ANOVA test was performed (p -value< 0.001). This resulted in three sets of up- and down-regulated genes; intersecting these allowed discriminating the differentially expressed genes (DEGs) exclusive to each the cell population. Principal Component Analysis (PCA) was then performed using the 24 samples as observations and 2770 genes (the total number of DEGs) as variables. To identify DEGs in control and treated populations, we performed 1-way ANOVA test using high p-value cutoff (0.05). A PCA analysis using 21 samples as observations (3 samples with PC1 score values ≈0 were excluded) and 370 DEGs indicated a clear-cut discrimination according to treatment with antibody. To identify DEGs between treated samples and controls, considering those DEGs among phenotypes, a 2-way ANOVA test was performed (p-value < 0.05). Gene Ontology categories corresponding to the 6 resulting genes sets were interrogated using the DAVID Database (Huang et al. 2007) for the functional enrichment analysis. Only functional categories from Biological Processes annotations were considered, filtering out those that involved less than 10 genes.

### Single cell RNA Seq

Cells were sorted in 2.3 µl of lysis buffer containing 0.2% Triton X-100 (Sigma-Aldrich) and 1U of Superase-In RNase Inhibitor (Ambion) and processed following the Smart-Seq2 protocol (^26^). Following preamplification, all samples were purified using Ampure XP beads (Beckman Coulter) at a ratio of 1:0.6. The cDNA was then quantified using the Quant-iT PicoGreen dsDNA Assay Kit (Thermo Fisher) and size distributions were checked on high-sensitivity DNA chips (Agilent Bioanalyzer). Samples were used to construct Nextera XT libraries (Illumina) from 125 pg of preamplified cDNA. Libraries were purified and size selected (0.5X-0.7X) using Ampure XP beads. Then, libraries were quantified using KAPA qPCR quantification kit (KAPA Biosystems) and sequenced in a Illumina HiSeq 4000 instrument.

### Bioinformatic Analysis of sc RNA-seq

Reads were mapped to the *Mus musculus* genome (EMSEMBL GRCm38.p4 Release 81) and ERCC sequences using *GSNAP* (version 2014-10-07) with -B 5 (batch mode 5) -n 1 (maximum pathsallowed: 1) -Q (if maximum paths more than n, not print) -N 1 (look for novel splicing). *HTseq-count* was used to count reads mapped to each gene with -s no (non-strand specific mode) ^38^ For further analyses, we only retained samples that had (1) more than 100,000 reads mapped to nuclear mRNAs; (2) a proportion of reads mapped to genes (nuclear + mitochondrial) relative to total reads above 20%; and (3) less than 20% of mapped reads allocated to mitochondrial genes. Overall, 483 cells (84%) passed our quality controls distributed as 239 cells from 4 control embryos and 244 cells from 4 a-Dll4 treated embryos.

Data were normalised for sequencing depth and RNA quantity using size factors calculated on endogenous genes ^39^]). Highly variable genes were identified as described^39^), using a false discovery rate threshold equal to 0.1. Only highly variable genes were considered to perform tSNE analysis, using the *Rtsne* package from *R*.

Clusters were identified using *ICGS* in *AltAnalyze* package ^25^. For integration with previously published dataset ^26^, the data was downloaded from GEO repository (GSE67120) and processed as indicated above. Differentially expressed genes were obtained for endothelial cells, pre-HSCs (T1 and T2), HSCs (E12 and E14) by comparing each group with the rest of the dataset. The union of the top 50 differentially expressed genes (log2 fold change > 2, adjusted p value (BH method) < 0.1 and baseMean > 50) from each comparison was selected to perform hierarchical clustering in our dataset using ‘ward.D2’ as the agglomeration method.

Cell cycle stage was predicted using the function *cyclone* in R package *scran* ^40^. Differential expression analysis was performed using the package DESeq2 (version 1.18.1) ^41^.

### Clonal analysis

VeCadherinCre^ERT^:R26R-Confetti analysis was performed using endogenous fluorescence of the transgene and additional staining for Kit or CD31. Distinction of the confetti colors was done under a confocal microscope using appropriate excitation length and filter combinations (Leica SP5 or SP8).

### Mathematical modelling

We use a markov chain model to simulate the formation of the cluster. The simulation has two stages: 1. Stage of cluster initiation. 2. Stage of cluster proliferation and recruitment of more cells from the endothelial tissue.

The simulation algorithm is as follows:

A time step loop is started from time 0 until some *T*_*life time*_ which we define as the cluster life time. This is the time after which a cluster is released from the aorta and can no longer be captured in the measurement. Assuming that cluster initiation is a random event, clusters can be initiated any time between 0 and *T*_*life time*_ such that the probability per unit time for cluster initiation is uniform and is given by 1/*T*_*life time*_. Since in physiological conditions all two-cell clusters we observed experimentally are either all colored or all non-colored we assume that the initiation stage always starts with two cells. After an initiation event has occurred the simulation moves on to the cluster proliferation and cell recruitment stage.

In the proliferation and cell recruitment stage there are two possible occurrences in each time step:

1. Each cell in the cluster can divide. Since we observed an average cell cycle time of 12 hours, we assumed that cell division can occur between 11-13 hours after the last division event. The probability of division increases linearly between 11-13 hours such that the probability of division at 10 hours after the last division is 0 and the probability of division 13 hours after the last division is 1.
2. In each time step a recruitment event can occur with probability per unit time of 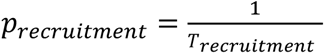 (with units of *hour*^−1^). The color of the recruited cell goes according to the percent of colored cells in the endothelial tissue such that the probability per unit time of recruitment of a colored cell is *p*_*recruitment colored*_ = *p*_*recruitment*_ * *percent of colored cells in the tissue* and the probability per unit time of recruitment of a non-colored cell is *p*_*recruitment colored*_ = *p*_*recruitment*_ * *percent of non – colored cells in the tissue*. After a cell is recruited, it is assumed that it has another cell cycle of 11-13 hours before the next division event.

There are two unknown parameters determining the formation of the cluster: *T*_*life time*_ and *p*_*recruitment*_. First, we determined what are the best *T*_*life time*_ and *p*_*recruitment*_ that fit the data of the IgG treated cells. We ran the simulation 4000 times for each combination of the two parameters. From the simulations of each set of parameters, we fitted the cluster size and ratio of colored cells in the cluster to the measured data (*Average experimental cluster size* = 5.5 ± 2.9; *Average experimental ratio of colored cells* = 0.54 ± 0.38). The fit was done using a Hotelling’s T2 test. The best fitting *T*_*life time*_ and *p*_*recruitment*_ we extracted are *T*_*life time*_ = 21, *p*_*recruitment*_ = 1/11.

Next, we determined the effect of α treated on *p*_*recruitment*_. The simulation for the Dll4 treated clusters is as follows:

1. Initially, the clusters develop without the effect of αDll4. The simulation is run according to the parameters determined in the IgG treated cells as described above. This is the stage of cluster formation before αDll4 treatment.
2. After a cluster is initiated, it continues to develop for a period of *T*_*life time*_, however this time it assumed that it is treated with αDll4 in the final 5 hours of the simulation. During the αDll4 treatment the probability per unit time of recruitment is assumed to be 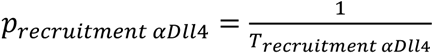

We determined the *p*_*recruitment αdll4*_ that fits the measured data of α-Dll4 treated clusters the best. We ran the simulation 4000 for each possible *p*_*recruitment αdll4*_, and again fitted the cluster size and ratio of colored cells in the cluster to the measured data *(Average experimental cluster size αDll4-treated cells= 7.1± 6.7; Average experimental ratio of colored cells=0.50± 0.37)*.

The Matlab code for the stochastic simulation is available at https://github.com/OhadGolan/ClusterFormation.

### Statistical analyses

Statistical comparison between two or more groups was performed using an unpaired, Student’s T-Test with equal variance. For the mathematical model the Hotelling’s T-Squared test multivariate counterpart of the T-Test was used ^42^. For comparing distribution of clusters in size categories the non-parametric Mann-Whitney U Test was applied.

## AUTHOR CONTRIBUTION

C.P. designed and performed most experiments, analyzed the data and wrote the manuscript, O.G, F.J.C.N, R.T. and CRH designed and performed experiments and analyzed the data. F.C., J.G., S.J.K., R.S., S.A.M. and A.M. performed experiments. X.W., Y.G. and PC performed bioinformatics analysis. E.D., L.E., B.G. and D.S. designed and supervised experiments and data and contributed with valuable tools. A.B. designed and supervised the research project, analyzed the data, and wrote the manuscript.

## ACKNOWLEDGEMENTS

We thank Thomas Graf (CRG), Minhong Yan (Genentech), Georges Lacaud (CRUK Manchester Institut) and Ralf Adams (Max Planck Institute for Molecular Medicine, Münster) for important mice models and reagents. Thanks to Catherine Robin, Timo Zimmermannand Raul Gómez-Riera for imaging advice. Thanks to the PRBB facilities: animal facility, flow cytometry, genomics, CNAG-CRG, confocal microscopy for critical technical assistance. Thanks to all the members of the lab for helpful critical discussions. This research was funded by the Ministerio de Economía y Competitividad (SAF2016 -75613-R) and Generalitat de Catalunya, Agència de Gestió d’Ajuds Universitaris i de Recerca (AGAUR) (2017 SGR 135) and PERIS-SLT002/16/00299 to AB. CP was a recipient of Juan de la Cierva fellowship (FJCI2014-19870), RT is a recipient of Beatriu de Pinos (2016 BP 00021), JG is a recipient of PERIS (SLT002/16/00070).

